# Human and generative AI integrate visual cues differently in a shape completion task

**DOI:** 10.64898/2025.12.17.694878

**Authors:** Jonathan Adams, Luca Serrière, Maximilian Jonathan Kothen, Maria Barbara Smorczewska, Filipp Schmidt, Yaniv Morgenstern

**Author notes:** shared first authors (Jonathan Adams, Luca Serrière). shared senior authors (Filipp Schmidt, Yaniv Morgenstern). Corresponding author information: Yaniv Morgenstern, Erasmus University Rotterdam Burgemeester Oudlaan 50, 3062 PA Rotterdam, The Netherlands Mail. Other author information: Yaniv Morgenstern, Mail, Filipp Schmidt, Mail, Maria Barbara Smorczewska, Mail, Maximilian Jonathan Kothen, Mail, Luca Serrière, Mail, Jonathan Adams, Mail.

## Abstract

In amodal completion observers perceive complete objects despite partial occlusion. When two object parts are divided by an occluder, completion can result in perceiving one or two objects. This phenomenon involves both lower-level cues (e.g., symmetry, contour continuity) and higher-level cues (e.g., prior knowledge). Experiment 1 investigates how occluder size, familiarity, and symmetry affect human completions using a drawing task. Narrow occluders and asymmetry promote single-shape completions, while familiarity and (global) symmetry promote two-shape interpretations. Good continuation emerges as the strongest cue, with symmetry and familiarity playing increasingly important roles as occluder width increases. Experiment 2 compares human performance with three state-of-the-art generative AI models. Models often generated creative but non-compliant outputs, altering even unoccluded regions. We restricted analysis to instruction-following generations, identified through ratings by naive observers. Among compliant outputs, models showed some human-like biases (e.g., more two-shape completions for wide occluders), but failed with higher-level cues. They did not use symmetry to guide completions and showed reversed familiarity effects. Our findings highlight differences between human and AI completions. Humans integrate low- and high-level cues, whereas compliant outputs from the AI models rely primarily on low-level pattern continuation. Current AI models lack the flexible integration of multiple representational levels that characterize human perception. This work establishes an analytical framework for evaluating whether next-generation models achieve more human-like visual reasoning.

**Highlights:**

- Humans see one or two objects behind occluders using geometric and semantic cues.
- Human drawings and generative AI completions show how different cues modulate perception
- Good continuation dominates human and AI shape completions
- Symmetry and familiarity guide humans, but not instruction-following AI outputs
- Humans integrate multiple levels; instruction-following AI engages lower-level-processing.

## 1. Introduction

In our everyday lives, we are constantly making inferences to help us make sense of what we see (Busey & Zaki, 2004; Pasupathy et al., 2018). A particularly prominent example of such a ubiquitous visual inference is the perception of partly occluded objects (Rock & Palmer, 1990; Pasupathy et al., 2018). In most of our environments, many objects in our field of view are covered by other objects so that only some parts of them are visible; yet we do not experience them as mere fragments. Rather, we perceive complete objects, as a result of our visual system’s inferences about their “most likely” shape, color or texture (Anderson & Julesz, 1995; Pasupathy et al., 2018).

This phenomenon, known as amodal shape completion (Gauker, 2024; Rock & Palmer, 1990; Van Lier & Ekroll, 2020; Mezey, 1992; Zhang & Lu, 2004), refers to the perceptual inference of whole objects despite partial occlusion. Even in the complexity of natural scenes, where countless completions are possible, humans often show remarkable consistency in how they infer or fill in occluded parts. For example, in **Figure 1A**, the partially hidden objects could be interpreted in many ways, such as disconnected front and back halves, as one elongated elephant, as two elephants with the rightmost one closer, as two elephants with the leftmost face closer, and so on. Yet, observers typically perceive them as two complete elephants, with the leftmost appearing nearer. Such cross-observer consistency in amodal completion reflects general purpose mechanisms in the visual system that overcome the ambiguities introduced by occlusion.

**Figure 1.**
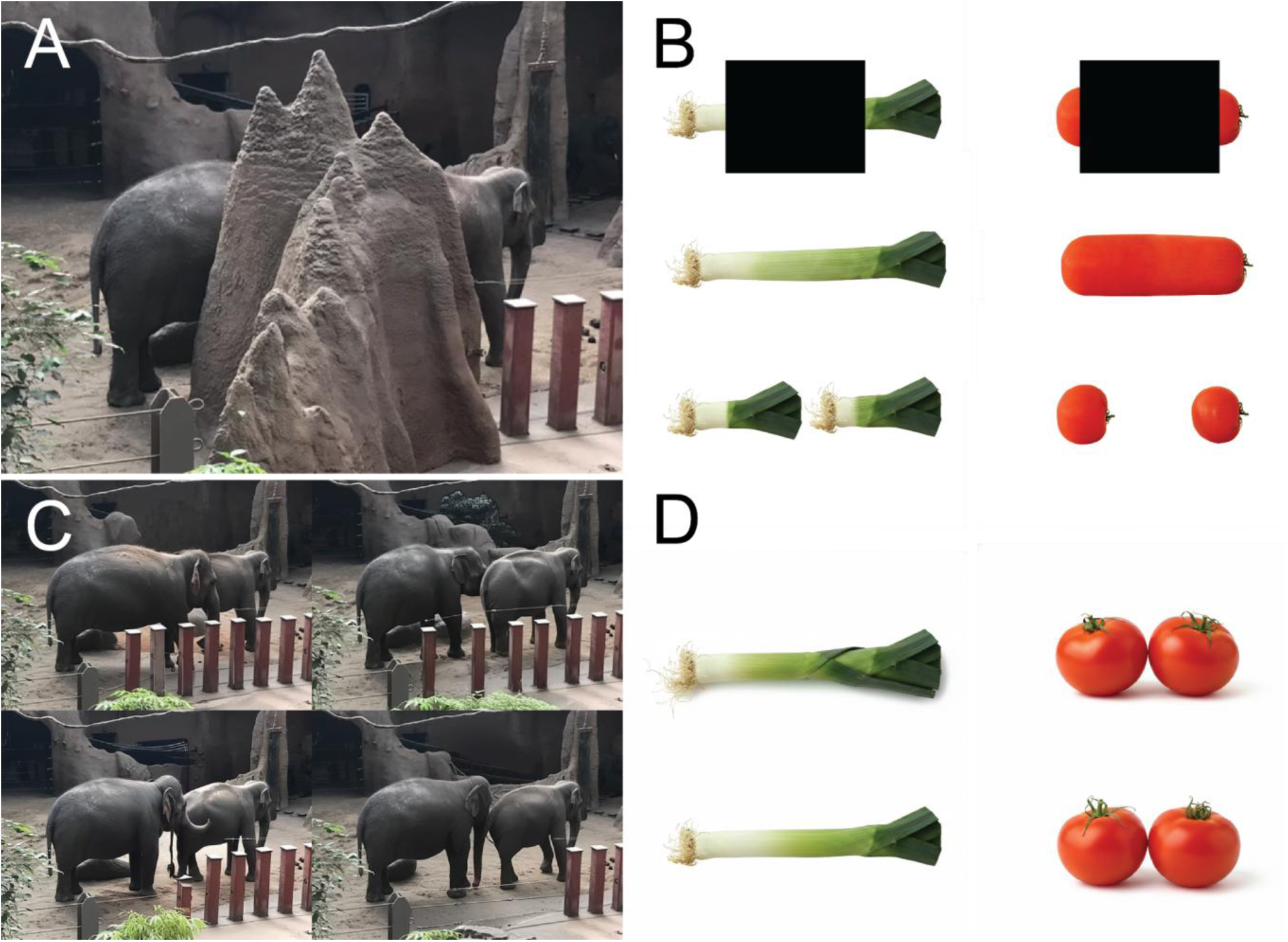
Examples for amodal completions into one or two shapes. (**A**) Elephants in the Blijdorp zoo in Rotterdam (photo copyright 2025, author Y. M.), and (**B**) convergent and divergent conditions in Yun et al., 2018. In convergent conditions, lower level cues (good continuation) and high-level cues (semantic knowledge about object size/shape) both support the same completion (e.g., the green onion’s elongated shape allows both smooth contour continuation and a plausible single-object interpretation. In divergent conditions, these cues conflict (e.g., the occluder is too wide for a single tomato to span the gag), so semantic knowledge suggests two objects despite the potential for good continuation (**C-D**) Example completion results for a state-of-the-art AI model (Google’s Gemini 2.5) for the occluded image section behind the (**C**) rock, or (**D**) black rectangular occluders.

The factors that influence this general purpose mechanism can be broadly divided into lower-level and higher-level cues (Thielen et al., 2024, Yun et al., 2018). (1) Lower-level cues are structural shape features such as regularity, complexity, symmetry or curvature (De Wit & van Lier, 2002, Domini & Caudek, 2003; Baker and Kellman, 2021). (2) Higher-level cues, in contrast, are context-dependent such as previous knowledge, semantics, or cognitive heuristics (Domini & Caudek, 2003, Norman et al., 2005). Consider, for example, the elephant(s) in **Figure 1A**: lower-level cues such as regularity and good continuation - the assumption that contours continue along their ‘natural path’ behind occluders (Wagemans et al., 2012; Wouterlood & Boselie, 1992) - might suggest that the visible head and rear are part of the same elephant. However, high-level cues, such as our knowledge about the typical shape and size of elephants, lead most observers to conclude that the photo is showing two partially occluded elephants.

In the real world, lower- and higher-level cues operate together, influencing perceptual inferences in tandem, both within, and across levels (Yun et al., 2018; Norman et al., 2005; Lappin et al., 2011). Kanizsa and colleagues (1975; 1979), for example, studied cases where two lower-level cues suggested different completions. They generated stimuli where good continuation cues suggested an asymmetric final shape, whereas symmetry favored a symmetric one. When the cues conflicted, they found the preference for good continuation generally outweighed the preference for symmetry.

Yun et al. (2018) on the other hand, studied cases where lower-level cues diverged from higher-level cues. Participants viewed partially occluded fruits - including either typically elongated (e.g. leek) or round (e.g. tomatoes) vegetables or fruits (**Figure 1B**, top row) - followed by continuous (**Figure 1B**, middle row) or discontinuous (**Figure 1B**, bottom row) versions of the same vegetables (also cf. Hazenberg & van Lier, 2016). The design allowed the authors to pit cue types against each other. For elongated vegetables such as the leek, low-level good continuation and higher-level knowledge cues converged, both suggesting a single leek stalk. For shorter vegetables like tomatoes, however, the cues diverged:good continuation favored a single tomato (unusually long), whereas knowledge about natural fruit shapes suggested the presence of two tomatoes. Yun et al. (2018) found that both cues influence shape completion, but the general tendency towards good continuation is overridden when it produces completions that violate object-specific knowledge.

Evidently, amodal completion involves the processing of complex and, at times, conflicting cues. In the case of disambiguating whether an occluded image contains one or two shapes, the overwhelming consistency in completions between participants suggests that the human visual system processes these cues in a systematic manner. Although this systematicity in amodal completion seems apparent in human vision, it is less clear in computer vision and artificial intelligence (Baker et al., 2023). In the context of computer vision, recent work has focused on amodal completion in realistic visual scenes. Different kinds of computer vision models have been tasked with amodally completing (1) shapes (e.g. contours) and (2) appearances (e.g. textures), as well as judging the (3) spatial order of objects in a scene (e.g. which objects are closer in depth) (see Figure 1 in Ao, et al., 2023). Although these three tasks fall under the same umbrella, they differ in both the information they require and the stage of processing they represent: shape completion involves inferring coherent object boundaries, appearance completion fills in surface properties such as color or texture, and spatial-order judgments require reasoning about depth and relative positioning. Shape completion is generally regarded as the foundational step, without clearly defined boundaries, appearance completion and spatial-order judgments may be ill-posed. For this reason, our work focuses on the first stage, enabling us to directly examine how models, like the human visual system, integrate lower- and higher-level cues when resolving contour ambiguities. Such efforts in computer vision have largely aimed at improving application performance, rather than explaining mechanisms of human vision.

Moreover, it remains unclear whether computer vision models can integrate lower- and higher-level cues in a manner comparable to the visual system. Recently, the emergence of large multimodal models (LMMs), which integrate semantic and visual information, raises the possibility that such models might incorporate both cue types and therefore achieve performance that more closely resembles human perception. These models represent a departure from traditional computer vision approaches: trained on vast datasets encompassing both visual and linguistic information, they can leverage semantic knowledge alongside perceptual features. Contemporary LMMs have demonstrated remarkable capabilities in replicating human-like performance across diverse cognitive tasks, including passing variants of the Turing test and achieving human-level scores on standardized reasoning benchmarks (Jones & Bergen, 2025). Moreover, machine learning models have shown striking correspondence with human behavior and neural representations, successfully predicting behavioral responses (Song et al., 2025) and mirroring hierarchical processing in human visual cortex (e.g., Yamins & DiCarlo, 2016; Schrimpf et al., 2020).

To illustrate this potential, consider the example shown in **Figure 1C**. When presented with an image of an elephant partially occluded by a rock (**Figure 1A**), similar to human observers a state-of-the-art LMM (Google’s Gemini) tends to generate two elephants. Similarly, a partly occluded leek stalk or two tomatoes are completed in line with the vegetables’ natural appearances (**Figure 1D**). This convergence suggests that, through exposure to a large corpora of natural images, these models acquire perceptual priors similar to those of human observers.

However, such demonstrations, while suggestive, leave critical questions unanswered. Previous research has shown that image-based neural networks can exhibit systematic biases that diverge from human perception (e.g., texture bias in object recognition networks; Geirhos et al., 2020), additionally it has been shown that whilst shape is one of the most important features for object classification in human vision (Pentland, 1986), models do not rely on shape for classification (Geirhos et al., 2018; Baker et al., 2018). Lastly, although multimodal models combine language and images, they seem to struggle with following task instructions, and often make odd additions clearly prohibited in the prompt. Consider for instance the tomatoes in **Figure 1D**, even though the model recognized the two tomatoes, it changed their orientation regardless of the instructions to not change anything that was outside of the occluder. These findings underscore that superficial agreement on specific examples does not guarantee that models process visual information in human-like ways. Furthermore, most studies demonstrating model-human correspondences have focused on general object recognition or categorization tasks. Far less is known about whether LMMs exhibit human-like behavior on specific perceptual phenomena such as amodal shape completion, particularly when systematically manipulating the very cues that humans are known to integrate.

### 1.1 Our study

The current study thus has two goals. In a first experiment, we investigated how lower-level cues, such as good continuation and symmetry, and higher-level knowledge influence whether visible parts are perceived as belonging to one or more objects (**Figure 1A**). To capture these percepts in an immediate and information-rich format, we used a drawing task (Fan et al., 2023). In a second experiment we aimed to assess whether three recent LMMs show similar performance on complex amodal completion tasks by comparing their outputs to the human data.

### 1.2 Hypotheses Experiment 1: Completion by humans

For Experiment 1, we examined how good continuation, familiarity, and global symmetry influence whether occluded shapes are perceived as one object or two.

#### Good continuation

We hypothesized that good continuation would significantly affect completion. Since contour extrapolation becomes more difficult over longer distances (Singh & Fulvio, 2005), we hypothesize that narrow occluders would lead to more one-object completions than wide occluders. However, based on Kanizsa and colleagues’ (1975, 1979) observations that good continuation can span large gaps (**Figure 2**), we still expected some one-object completions even in the wide occluder conditions.

**Figure 2.**
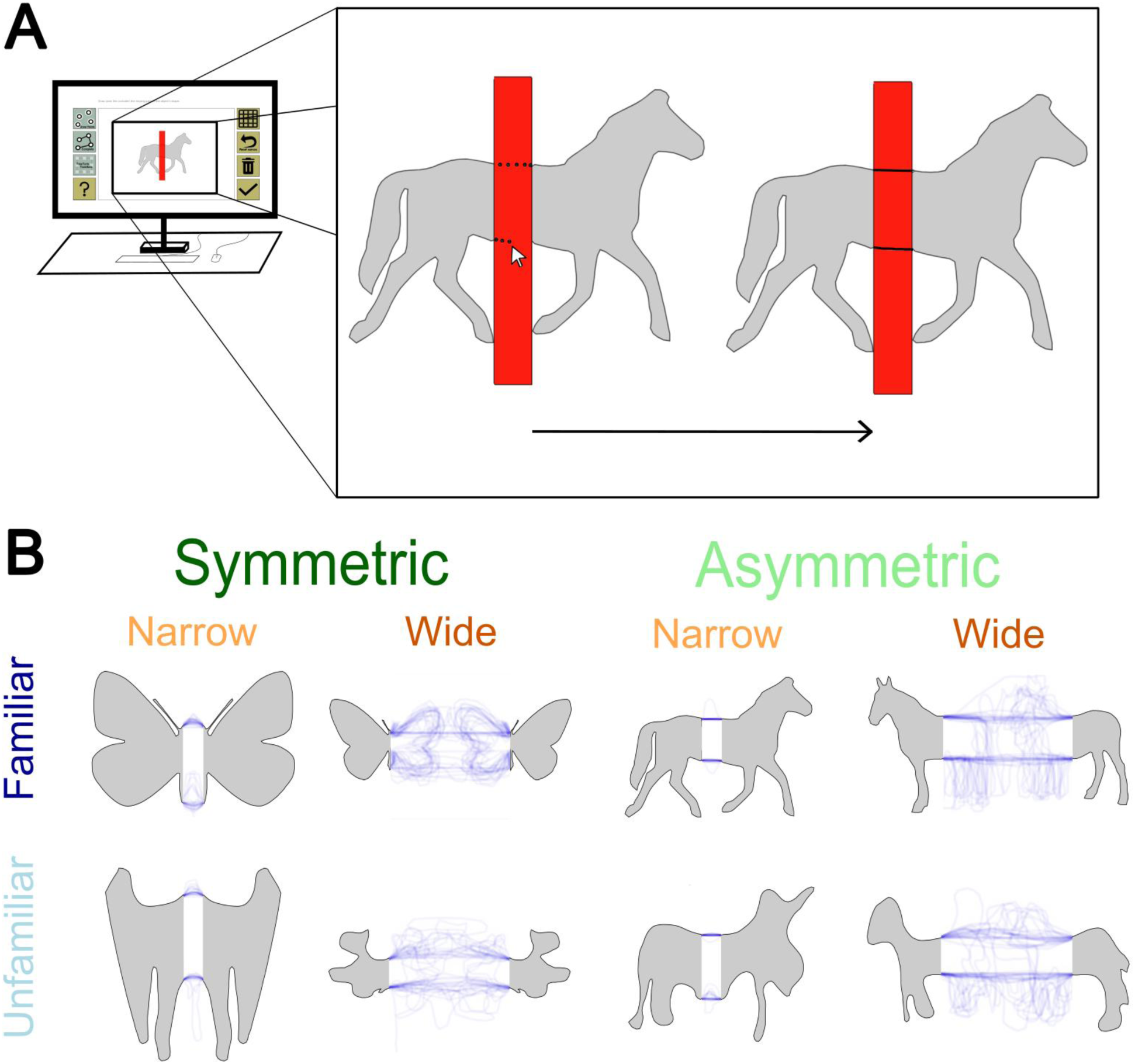
Experimental Paradigm for the Shape Completion Task. (**A**) Experimental interface showing a partially occluded shape with a red occluder and the digital drawing tools available to participants. The magnified inset demonstrates the progression of the completion process with two sequential states of the same horse stimulus, showing how participants could draw connecting lines across the occluder by placing dots and then using one of the available tools to connect them. (**B**) The eight experimental conditions created by crossing three variables: shape type (familiar vs. unfamiliar), symmetry (symmetric vs. asymmetric), and occluder size (narrow vs. wide). Blue traces represent individual drawings of participants from Experiment 1.

#### Familiarity

With wide occluders, we predicted that familiarity would modulate completion. For familiar shapes, completing them as one might yield implausible objects (e.g., an unnaturally long elephant), making two-object completions more likely. With narrow occluders, both familiar and unfamiliar shapes would retain plausible proportions when completed as one object (e.g., a normal-sized elephant), so no familiarity effect was expected.

#### Global symmetry

Global symmetry refers to axes of symmetry that involve all points of an image, here manipulated by mirroring one half of the shape to produce a single symmetry axis behind the occluder. Two opposing hypotheses were possible.

1. *Grouping hypothesis:* According to Gestalt principles, symmetric parts are more likely to be grouped together (Apthorp & Bell, 2014; Wagemans et al., 2012), and symmetry often signals a single object (Parovel & Vezzani, 2002). Thus, globally symmetric shapes should be more often completed as one.
2. *Axis-reinforcement hypothesis:* the presence of a global axis of symmetry increases sensitivity to possible local axes of symmetry (Nucci & Wagemans, 2007). In our stimuli, especially with wide occluders, visible information lies far from the global axis, which can reduce symmetry detection (Foster & Kahn, 1985). This might encourage the creation of closer local symmetry axes by completing shapes as two, as this would decrease the distance between points in the image and their relative axis of symmetry, which increases the overall ‘goodness’ of the array (Nucci & Wagemans, 2007).

### 1.3 Hypotheses Experiment 2: Completion by Large Multimodal Models

In Experiment 2 we tested three state-of-the-art large multimodel models (LMMs): Google’s Gemini 2.5 flash experimental image generation (Google, 2025), OpenAI GPT image generation (OpenAI, 2025) and xAI Grok 3 (xAI, 2025). These models were selected because, at the time of testing, they represented the latest LMM’s suitable for our task. We compared model outputs to human data at two levels. First, we investigated the tendency to complete shapes as one shape or two shapes. Second,we tested the effects of global symmetry on these completions. As this investigation was highly exploratory and no prior research had assessed LMM performance on amodal completion, we did not formulate specific hypotheses.

### 1.4 Conclusion

This study extends on previous work in at least three ways. (1) Experiment 1 increases ecological validity by using complex, natural shapes to investigate the interaction between good continuation and symmetry. Additionally, both unfamiliar (abstract) and familiar (animal) shapes were selected to specifically study the effect of higher order knowledge cues. (2) Using a digital drawing paradigm allowed participants to report their completions directly, without being constrained by predefined interpretation options (Fan et al., 2023). (3) Finally, to our knowledge, Experiment 2 represents the first direct comparison between LMMs and humans in the context of amodal shape completion.

## 2. Methods

### 2.1 Open practices statement

All data and stimuli will be published via https://zenodo.org/ after publication. Data were analyzed using Matlab 2023ba, version 9.4.0 (Natick, MA: The MathWorks Inc., 2018). This study’s design and its analysis were not pre-registered.

### 2.2 Participants and procedures

The experiments were approved by the Local Ethics Committees of the Department of Psychology and Sports Sciences of the Justus Liebig University Giessen (LEK-2017-0046) and Erasmus University Rotterdam (ETH2425-0766). Participants were recruited via university mailing lists. For Experiment 1, data collection took place at the Erasmus Behavioural Lab at Erasmus University Rotterdam. Participants in all experiments had normal or corrected-to-normal vision and were naïve to the purpose of the study. The sample consisted of *N* = 29 participants (24 female, 5 male), with a mean age of *M* = 20.5 years (*SD* = 2.1, range = 18–29). Participants gave informed consent and were treated in accordance with the ethical guidelines of the American Psychological Association (APA). All participants received financial compensation or course credits for their participation.

At the beginning of the experiment, participants first read the consent form, and their continuance was interpreted as consent (Justus Liebig University Giessen: participants signed the form). Participants were also provided with a physical sheet containing instructions and explanations for the tools and user interface. Demographic information was collected at the start of the experiment, with participants selecting their gender (male, female, or other) and indicating their age.

### 2.3 Experiment 1: Completion by humans

#### 2.3.1 Design

The experiment employed a 2 × 2 × 2 within-subjects design with three categorical variables:

1. **Shape** Familiarity: Familiar (animal) vs. Unfamiliar (abstract) shapes,
2. **Symmetry**: Symmetric vs. Asymmetric shapes, and
3. **Occluder Size**: Narrow vs. Wide occluders.

These variables created eight distinct experimental conditions, as illustrated in **Figure 2B**. The stimuli across the narrow and wide occluder conditions were semantically matched to ensure that for each familiar object (e.g., a narrowly occluded animal like a butterfly), there were corresponding stimuli in the other conditions (e.g., a widely occluded butterfly).

#### 2.3.2 Materials

The stimuli represented either familiar animal shapes (e.g., bear, butterfly) or unfamiliar abstract shapes. The animal shapes were sourced from PhyloPic.org, and the abstract shapes were generated using a Generative Adversarial Network (GAN) designed to produce animal-like forms (from Morgenstern et al., 2021).

The occluders varied in size, affecting the distance between the visible parts of the shape. In the "Wide" occluder condition, the occluder was sufficiently broad to leave substantial space between the visible parts, allowing the possibility of interpreting them as two separate objects. In the "Narrow" condition, the occluder was thin enough that completing the shape as two independent objects was less plausible.

The experimental task was implemented using JavaScript, jsPsych, and HTML, creating a full-screen interactive digital drawing environment. The custom-built interface allowed participants to draw and manipulate shapes with precision using various tools and functionalities.

#### 2.3.3 Computer Interface and Tools

The computer application, built upon the framework developed by Van Geert et al., (2025), provided participants with multiple interactive tools for the drawing task (**Figure 2A**):

1. **Draw Points:** The default tool allows participants to place dots at the cursor location by pressing the left mouse button on the canvas.
2. **Join Points:** Connected all placed points in the sequence they were placed, creating line segments.
3. **Free Transform:** Enabled adjustment of spatial positions of the drawn points and complete rotations of the entire drawing.
4. **Grid:** Provided an optional grid overlay to assist with precision in drawing.
5. **Reset Canvas:** Removed all drawn elements from the canvas.
6. **Delete Object:** Allowed for deletion of selected dots or lines.
7. **Zoom:** Participants could zoom in or out on the canvas using the mouse wheel for finer detail work.
8. **Done:** Finalized the current drawing and advanced to the next stimulus.
9. **Help (?):** Provided access to review instructions.

#### 2.3.4 Experimental Setup

Participants completed the experiment in individual, sound-reducing cubicles. Each station was equipped with a computer monitor, keyboard, and mouse. Participants were positioned at an arm’s length distance (approximately 50-60 cm) from the monitor to ensure standardized viewing conditions across all subjects. Experimenters were available to provide assistance if participants had questions or encountered difficulties with the interface.

#### 2.3.5 Procedure

Before beginning the experimental trials, all participants completed a comprehensive tutorial that explained each function of the interface through a series of instructional screens, followed by a practice stimulus to familiarize them with the task environment. The experimental task required participants to complete partially occluded shapes by drawing connecting lines between visible parts. Participants were explicitly instructed that "completing" meant connecting the visible unfinished edges to form a coherent, closed shape, which could be accomplished either by connecting edges across the occluder (completing as one object) or by connecting neighboring unfinished edges (completing as two separate objects). The only requirement was that the completion formed a coherent shape, allowing for multiple possible solutions.

For each stimulus, participants placed dots at key points along the visible edges, connected these points using the "Join Points" function, and refined their drawings using the various tools until satisfied with their completion. The experimental session presented 40 different occluded shapes in randomized order, representing all combinations of shape familiarity, symmetry, and occluder width conditions. Sessions typically lasted between 30-45 minutes depending on individual participant pace. Upon completion, participants received a debriefing form explaining the experiment’s purpose and scientific approach.

#### 2.3.5 Analysis

##### Screening

A total of 1,160 stimuli (40 drawings from each of 29 participants) were collected in Experiment 1. However, some of the participant drawings did not adhere to the instructions. An initial screening conducted by 5 of the 6 authors identified 283 drawings to be ambiguous for the following reasons: (1) one-sided closures (35 drawings) - partially completed drawing that were only closed on one side; (2) incomplete extensions (92 drawings) - drawings that did not reach the occluder’s edge; (3) artifact lines (137 drawings) - drawings with extra lines that appeared as noise or errors; (4) beyond-occluder drawings (12 drawings) - drawings that were exclusively outside the occluder; and (5) no drawings (7 drawings). Examples and more detailed description of each category are provided in **Supplementary Material S1**. Categorisation was determined by majority vote based on judgements from 5 of the 6 authors.

From these erroneous drawings, we attempted to recover some of 137 samples from ‘artifact lines’ in category (3). Using a length-based heuristic, we edited single contours that were greater than 3 points, applying this only to points spanning at least 10% of the occluder edge’s largest length. This heuristic identified and removed lines with atypical lengths that were likely drawn in error. As a result, we recovered 109 drawings from the 137 samples with ‘artifact lines’, yielding an updated dataset of 986 stimuli (85% of the original set).

##### Classifying Drawings as One or Two

5 participants (mean age = 26.0 years, *SD* =2.5, range = 21 - 29; 3 Female, 2 Male) were recruited to rate the drawings created in the experiment. The raters were students associated with the respective research teams from *Erasmus University of Rotterdam* and *Justus Liebig University Giessen*, and were blind to the aims and scope of the experiment. The raters were instructed to rate the drawings as one of the following options: (1) a single continuous shape, (2) two distinct shapes or (3) neither option/not clear. The mode of the ratings was taken from the raters judgements. A total of 13 drawings (1%) were deemed unclear and were therefore excluded from our definitive analysis, leaving us with a final dataset of 974 rated stimuli.

##### Global vs. Local Symmetry Analysis

The drawings determined to have been completed as two distinct shapes were then rated to determine the presence of local and global axes of symmetry. The same five raters of the previous rating procedure were asked to determine if each drawing was drawn with the intention of creating local or global axes of symmetry. The instructions and further description of local and global axes of symmetries are detailed in **Supplementary Material S2**. The mode of these ratings was then taken to analyse whether the presence of global axes of symmetry in symmetric stimuli would induce a completion with local axes of symmetry.

### 2.4 Experiment 2: Completion by Large Multimodal Models

#### 2.4.1 Design

#### 2.4.2 Materials

In the second experiment, conducted in June 2025, we aimed to explore whether contemporary Large Multimodal Models (LMMs) exhibit human-like tendencies in amodal completion tasks. We compared the outputs of three state-of-the-art LMMs with the human drawings from Experiment 1. The models selected were OpenAI’s GPT-4o, Google’s Gemini 2.5 Flash, and xAI’s Grok, chosen as they represented the latest, most capable, and publicly accessible image generation technologies at the time the experiment was conducted. All programmatic interactions with OpenAI’s GPT-4o and Google’s Gemini Flash were executed using Python 3.12. The responses for xAI’s Grok were executed through the web browser as an API was not available at the time of writing.

The general procedure involved presenting each model with the same 40 occluded shape stimuli used in the human drawing task. To ensure a consistent and constrained task across all three platforms, a specific instructional prompt was engineered. This prompt, used for every stimulus, was designed to explicitly instruct the models to perform an inpainting task, focusing solely on completing the black silhouette without altering the visible portions of the image or the background. It read: *"This is an image of a black silhouette on a light gray background, partially covered by a dark grey quadrilateral occluder. Remove the occluder and fill in the occluded region in a natural way that completes the silhouette. Do not modify any part of the silhouette that is visible outside the occluder. That is, everything outside the occlusion must remain pixel-identical to the original image. Only replace the occluded region with pixels that plausibly belong to the silhouette, maintaining the overall visual consistency and realism of the figure."* Note that while human participants completed the task with red occluders, we used grey occluders as input for the models. Qualitative pilot testing indicated that grey occluders produced outputs that better adhered to task instructions, though this was not formally quantified. Although we tested several equivalent phrasings during pilot runs, qualitative results were similar across formulations, and we selected this version for clarity and reproducibility. In practice, models sometimes failed to follow the prompt precisely (e.g., altering visible regions or failing to remove the occluder). To account for this, we had independent human raters score each output for instruction adherence on a [100]-point scale, and analyses were restricted to outputs rated as adequately following the task (see details in 3.2.1).

The specific procedure for each model was tailored to its available interface. Interaction with OpenAI’s GPT-4o was automated via a Python script utilizing the *openai* library (version 1.35.7). We used the *client.images.edit* function with the *gpt-image-1* model, providing the 1024x1024 pixel stimulus and a corresponding binary mask to generate five completions. For Google’s Gemini 2.5 Flash, a Python script using the *google-generativeai* library (version 0.7.1) was employed to interact with the experimental *gemini-2.5-flash-exp-image-generation* model. The model was prompted to generate five unique variations for each stimulus, with the *temperature* parameter set to *0* to favor deterministic outputs. All other parameters, such as safety settings, were left to their API defaults. As a public API for xAI’s Grok was not available, a manual procedure was implemented using its web-based chat interface. Six human operators were given a standardized instruction sheet and tasked with processing the stimuli. This manual process yielded six completions per stimulus. To ensure a balanced comparison with the other models, five of these six Grok generations were randomly selected for each stimulus using the *random.sample* function from Python’s *random* library.

This process resulted in five completed variations for each of the 40 stimuli from each of the three models, creating a dataset of 200 images per model, totalling 600 images. The images were then prepared for human rating. To standardize the data and blind the raters to the generating model, all 600 files were converted to a 1024x1024 pixel PNG format and subsequently renamed with a unique numerical identifier from 1 to 600.

#### 2.4.3 Analysis

The anonymized images were subjected to the same rating and analysis pipelines as described in section 2.3.5. This allowed the four of the same five human raters and 1 new rater (mean age = 23.4 years, *SD* = 3.8, range = 19 - 29; 4 female, 1 male) to assess each model-generated completion for whether it represented one or two objects and for the presence of local and global symmetry, enabling a direct and unbiased comparison between human and AI performance.

## 3. Results

### 3.1 Experiment 1: Completion by humans

#### 3.1.1 Overview

**Figure 3** provides an overview of all drawings produced across the experimental conditions. Each row corresponds to object familiarity (top: familiar, bottom: unfamiliar), while columns indicate symmetry (left: asymmetric, right: symmetric). Within each panel, different shades represent drawings from different observers, with blue indicating single-shape completions and red indicating two-shape completions. Black occluders mark trials in which no valid drawing was classified, which occurred more frequently for narrow occluders (right side of each panel) than for wide occluders (left side of each panel).

**Figure 3.**
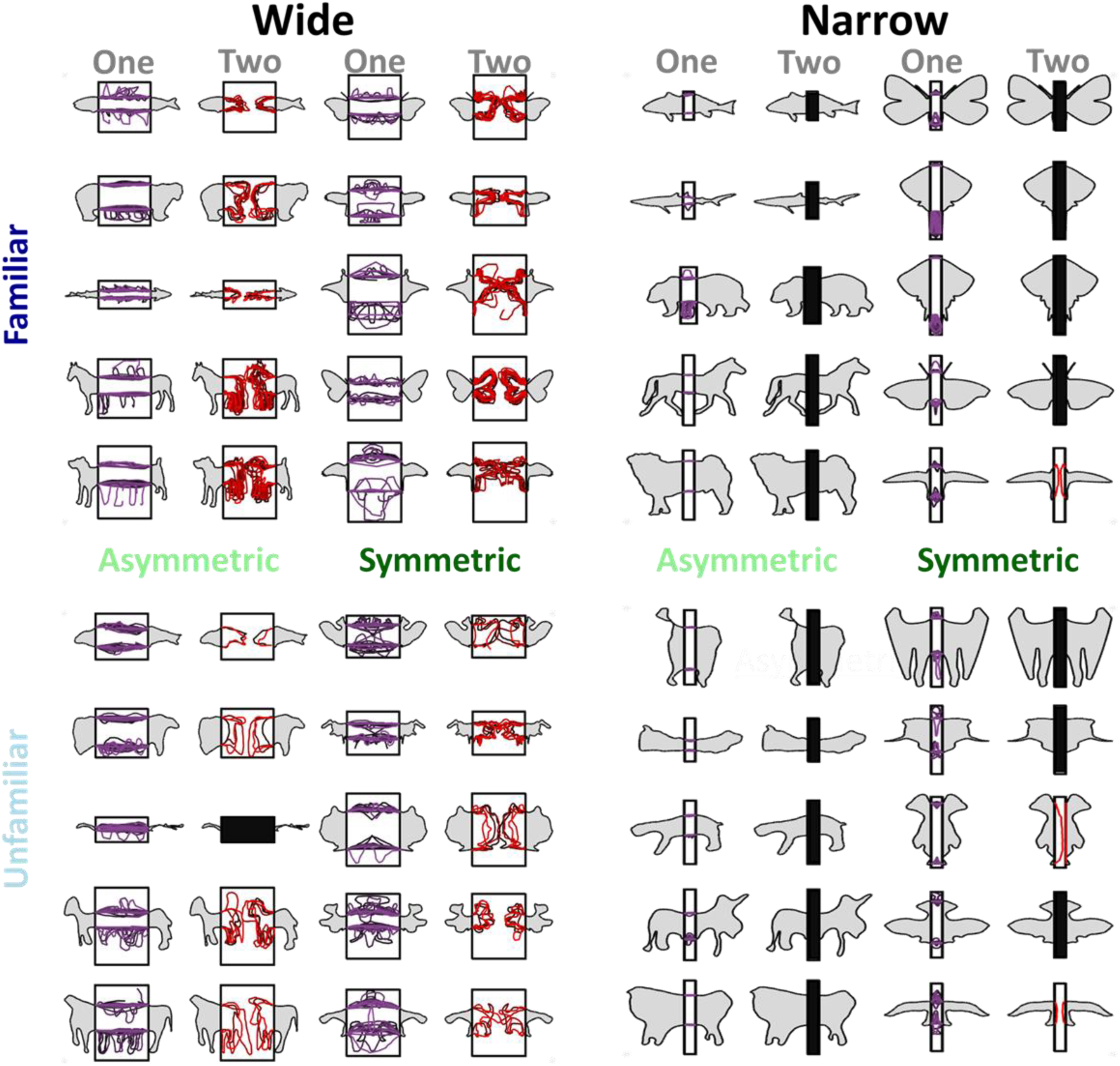
Dataset of human drawings across all experimental conditions. Different shades represent drawings by different observers with blue and red indicating single- and two-shape completions, respectively. Drawings were classified as containing one or two shapes based on the mode of five independent, naive raters. Black occluders denote conditions in which no drawing was classified, a pattern that occurred more frequently for two-shape completions in narrow (right) than wide (left) occluder conditions. Rows show familiar (top) and unfamiliar (bottom) objects; columns show asymmetric (left) and symmetric (right) alignments. Two-shape completions were most common for wide familiar objects and occurred more often in wide than narrow occluder conditions.

Several broad patterns are evident in this aggregated view. First, the frequency of two-shape completions (red) was higher for wide occluders than for narrow occluders, with the effect being most pronounced for familiar shapes. Second, narrow occluders tended to lead to consistent single-shape completions across participants, whereas wide occluders produced greater variability in both completion type and drawing form. Together, these observations hint at a strong role for good continuation in guiding completion: when the occluder is narrow, the implied contours align more clearly, supporting a single-shape interpretation; when wide, continuation is more difficult, leading to greater variability. This hypothesis is tested directly in the following analysis of occluder size (Section 3.1.2).

#### 3.1.2 Occluder Size

Building on the patterns observed in **Figure 3**, where narrow occluders elicited predominantly single-shape completions with low variability, we formally tested the influence of occluder size on completion behavior. Narrow occluders provided strong cues for good continuation, making it easier for participants to infer the hidden contours. In contrast, wide occluders disrupted these cues, leading to greater variability in the perceived completion (**Figures 3** and **4A**). Accordingly, the proportion of single-shape completions was significantly higher in the narrow condition than in the wide condition (Z= -12.75, *p*<.001). Only three drawings were completed as two distinct shapes in the narrow condition, demonstrating the near-uniformity of perception with strong good continuation cues. The wide occluder, by comparison, produced a more diverse set of completions, consistent with the weaker alignment of visible contours.

**Figure 4.**
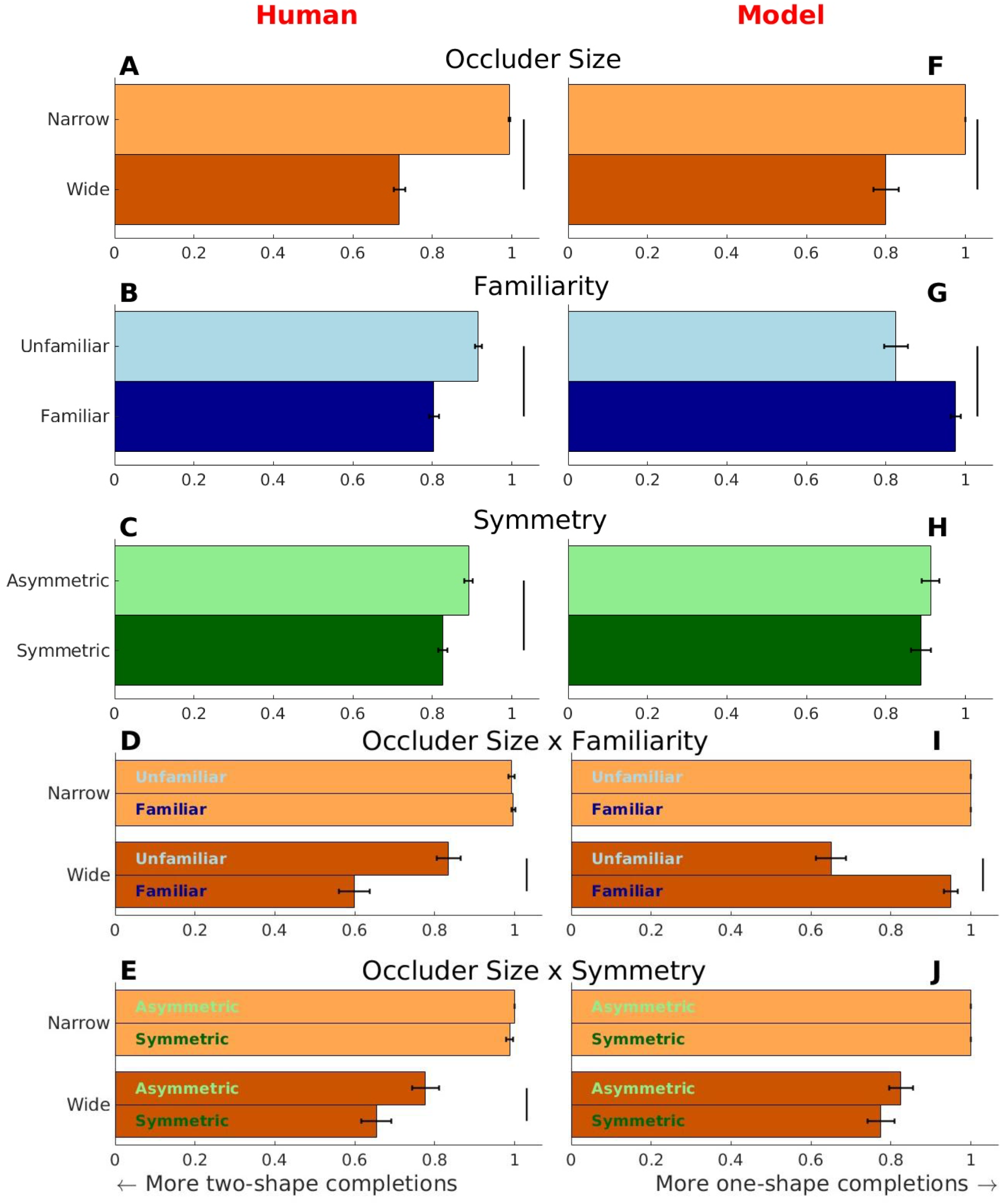
Results of Experiment 1 and 2 per condition. Proportions of shapes completed as one or two shapes depending on (**A** and **F**) occluder size (narrow, wide), (**B** and **G**) familiarity (unfamiliar, familiar), (**C** and **H**) symmetry (asymmetric, symmetric), a visualization of the interaction effect of (**D** and **I**) occluder size and familiarity, and (**E** and **J**) occluder size and symmetry. Vertical lines represent significant differences between conditions. Error bars represent the standard error of the mean. The left and right show the human and model shape completion tendencies respectively. All significance levels are Bonferroni corrected.

#### 3.1.3 Familiarity

We next examined whether prior knowledge about an object’s identity influenced completion, comparing familiar (animal) and unfamiliar (abstract) shapes. Overall, unfamiliar shapes were more likely to be completed as a single shape than familiar shapes (Z = -4.75, *p*<.001; **Figure 4B**). This suggests that when shape identity is unknown, participants may rely more heavily on the visible contour geometry, leading to a more uniform single-shape interpretation. In contrast, familiar shapes may activate stored representations that sometimes support alternative completions, increasing variability in responses. Note, that this difference between familiar and unfamiliar completions is only evident in the wide occluder condition (**Figure 4D**). In the narrow occluder condition, good continuation and familiarity converge: the strong continuation cues favor single-shape completions, and the narrow gap is too small to support recognizable two-shape configurations.

#### 3.1.4 Symmetry

We then investigated the influence of shape symmetry on completion outcomes. While previous literature presents mixed findings regarding symmetry’s role in perceptual grouping (Apthorp & Bell, 2014; Foster & Kahn, 1985; Nucci & Wagemans, 2007; Parovel & Vezzani, 2002; Wagemans et al., 2012), our data reveal a clear effect: asymmetric shapes were completed as single shapes more often than symmetric ones (Z = –2.89, *p* = .003; **Figure 4C**). This indicates that asymmetry may strengthen the perceptual grouping of occluded parts into a single object, whereas symmetry may encourage interpretations involving multiple parts. Just as with familiarity this effect is more prominently present in the wide occluder condition (**Figure 4E**).

#### 3.1.5 Global and Local Symmetries in Shapes Completed as Two

Finally, we analysed the symmetry structure of the N = 137 drawings completed as two separate shapes. Using the procedure described in the task instructions (section 2.3.5; See also **Supplementary Material S2**), five raters judged each image for the presence of global symmetry (across both shapes) and local symmetry (within an individual shape).

Drawings were coded as locally symmetric if at least one shape in the pair contained a single local axis of symmetry. Of the 137 two-object drawings, 56 were asymmetric. Among these, cases of “local-only” symmetry—where local symmetry was present in the absence of a global axis of symmetry—were extremely rare, occurring only twice (**Figure 5**). In all other instances local symmetry co-occurred with a global axis of symmetry. This pattern appeared more frequently overall, occurring in 77% of the familiar two-drawing trials, where prior expectations about typical object size or shape may play a role, and in 46% of the unfamiliar two-drawing trials, showing that the effect also emerges quite regularly when global symmetry is present even without such prior knowledge. The occurrence of local symmetry in globally symmetric stimuli was significant (Z = 5.79, *p <* 0.001). These results show an increased tendency to complete a symmetric stimulus adhering to locally symmetric patterns.

**Figure 5.**
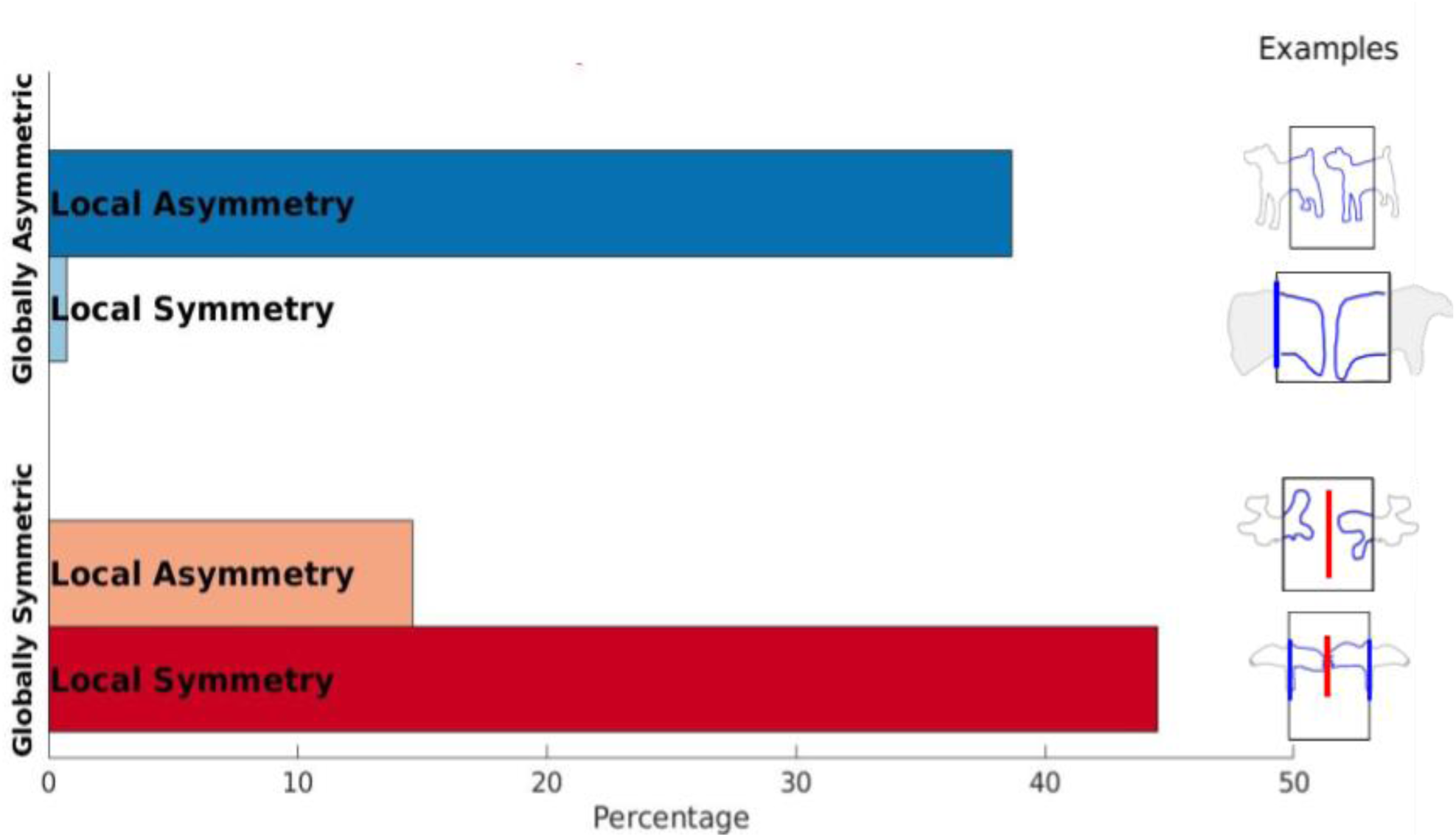
Local Symmetry Occurrences in Shapes Completed as Two in Human Data. The proportion of stimuli completed by humans with local symmetry in globally symmetric and asymmetric stimuli.

### 3.2 Experiment 2: Completion by Large Multimodal Models

#### 3.2.1 Overview

An overview of the images generated by the models is shown in **Figure 6**. Model outputs followed a continuum between non-compliant, creative outputs that violated task instructions, and compliant outputs that relied heavily on lower-level visual continuation. To quantify instruction adherence and identify meaningful outputs to compare AI and human completions, five independent observers rated all 600 generated images on a 0-100 scale for how well they followed instruction.

**Figure 6.**
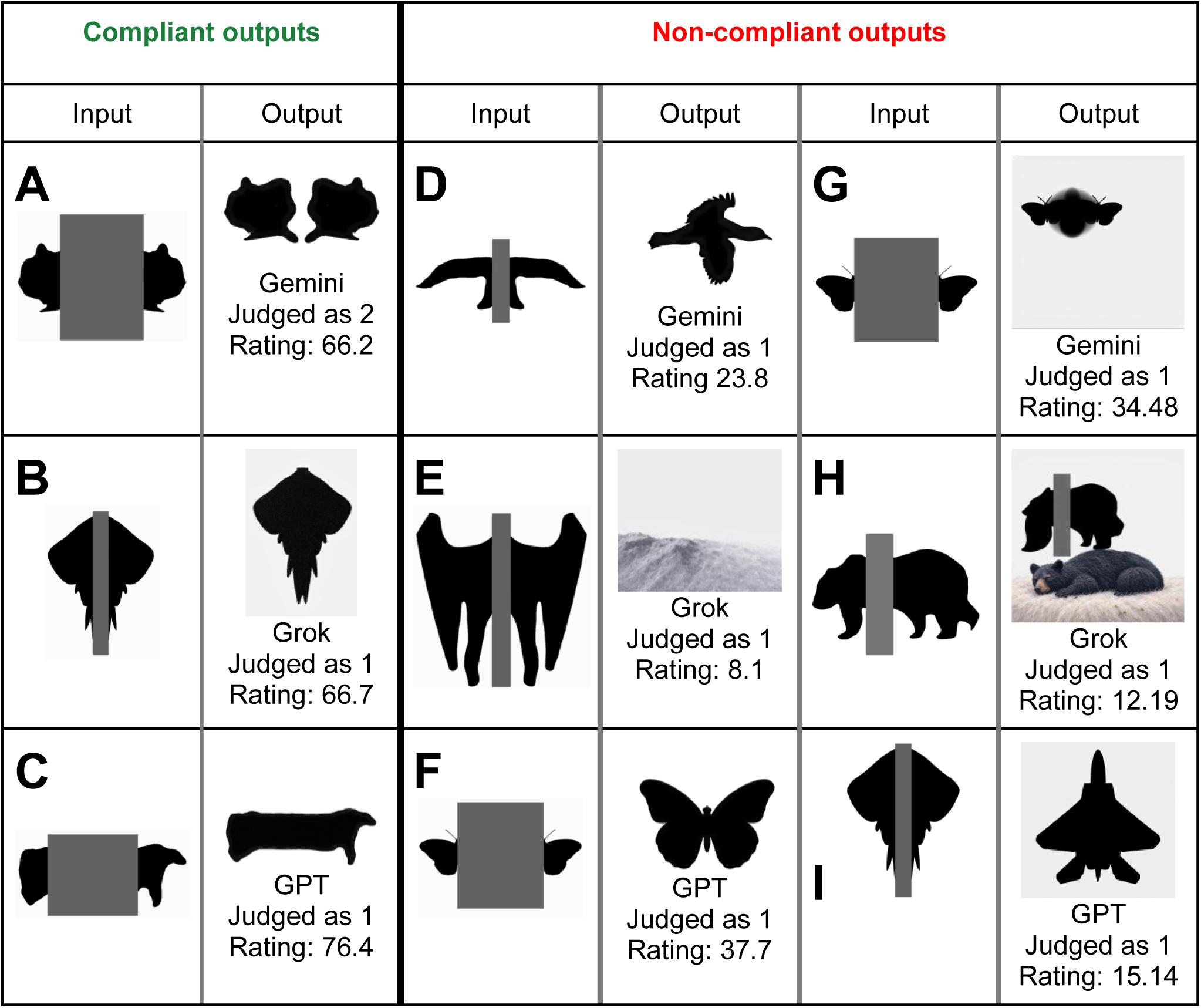
Examples of Compliant and Non-Compliant Model Outputs. Model completions showing compliant outputs that preserve unoccluded regions (**A,B,C**) and non-compliant outputs which were creative and did not follow instructions (**D-I**). Ratings show average compliance score (across 21 participants).

##### Creative but non-compliant outputs

A substantial proportion of the model outputs showed creative reinterpretations that failed to follow instructions (e.g., **Figure 6D-I**); a limited number of model-generated images achieved high instruction-following scores, with 344 out of 600 stimuli (57%) scoring below an average rating of 50. For example, prompts included the instruction to not change pixels outside of the rectangular occluder, but the outputs did often include such changes. This sometimes changed the properties we were manipulating by, for example, moving the unoccluded shapes closer (effectively reducing the occluder size, as shown in **Figure 6F**), or even removing all shapes and creating new images with no resemblance to the original stimulus (examples of such behaviour can be seen in **Figure 6D** and **6E**). At times, these creative deviations reflected high-level semantic interpretations: in **Figure 6D**, the model appears to have interpreted the novel shape as a bird viewed from a different vantage point and added wing-like features, while in **Figure 6F**, the model interpreted the symmetric wide-occluder stimulus as a single butterfly, effectively bringing the two parts together into one coherent object. The models frequently altered pixels outside the occluder region moving unoccluded shapes closer together or further apart showing their sensitivity to generating images that make sense visually, but violating the explicit constraint to preserve visible regions. These results underscore the LMMs’ tendency toward creative reinterpretations rather than faithful instruction-following, even when those instructions are explicitly stated in the prompts.

##### Compliant outputs

To meaningfully compare AI and human completions, we isolated outputs that adhered to task constraints. We analyzed only the highest-rated 20 images per condition (threshold: mean rating >50, balanced across eight conditions), ensuring comparison with outputs that respected the experimental manipulations. Critically, these instruction-following outputs showed a markedly different pattern: they relied primarily on local contour continuation and failed to incorporate high-level constraints like symmetry or semantic knowledge (see analyses below).

#### 3.2.2 Occluder Size

Among compliant outputs, occluder size plays a significant role in shape completion (Z = 4.22, *p* <0.001), similar to human completions. **Figure 4F** shows that narrow occluders are consistently completed as one shape with very little variability. Although this finding is similar to what is observed in human completions, the models would sometimes edit the images to bring the visible shapes closer or further apart. This violated instructions and introduced a stronger cue for good continuation, thereby distorting our experimental manipulation.

#### 3.2.3 Familiarity

Compliant outputs showed an effect of familiarity opposite to humans: familiar stimuli being significantly more likely to be completed as one shape compared to unfamiliar ones (Z = 3.16, *p* = 0.002, **Figure 4G**), with the effect concentrated in the wide occluder condition (**Figure 4I**). This pattern suggests that, unlike humans who prioritize familiarity as a strong cue for unity, the LMMs do not rely strictly on familiarity to guide their completions. Instead, they appear to favor good continuation, connecting shapes with smooth contours, regardless of whether the resulting completion turns out to be a familiar or unfamiliar outcome.

#### 3.2.4 Symmetry

Compliant outputs showed no significant effect of symmetry on model completions (Z = -0.53, p = 0.598, **Figure 4H**), including when analyzing only the wide occluder conditions where symmetry strongly influenced human completions (Z = -0.04, p = 0.968, **Figure 4J**). This absence is particularly revealing: while non-compliant outputs demonstrated sensitivity to global shape properties and semantic coherence (**Figure 6F**), compliant outputs failed to use symmetry to guide completion. This suggests that when constrained to follow instructions, models rely on local operations that cannot access or integrate global structural regularities.

#### 3.2.5 Global and Local Symmetries in Shapes Completed as Two

The model outputs were then rated by the same five observers as the human completions with respect to the occurrence of global and local axes of symmetry. While a total of 98 out of the 600 outputs were rated as being two distinct shapes. Only 11 of these met our threshold for compliant or instruction-following model output. Due to the lack of meaningful data the analyses for local and global symmetry completions are detailed in **Supplementary Materials S3**.

## 4. Discussion

Here, we investigated amodal shape completion, the continuation of shape contours behind an occluder, by asking human participants and state-of-the-art generative AI models to draw in the occluded sections. We use a generative task to study the specific case of amodal completion that can lead to qualitatively different results, namely completion of the visible parts as either one or two objects. In our first experiment we manipulated occluder size, symmetry, and familiarity to study the effect of different low- and higher-level cues on shape completion. Specifically, we analyzed participant’s drawings to test how good continuation, global symmetry and familiarity contributed to seeing visible parts of occluded objects as belonging to one continuous shape, or two separate shapes.

(1) To look at the effects of good continuation, we manipulated the size of the occluder (i.e. the distance between the two visible parts). We hypothesized that narrow occluders should result in stronger contour alignment across occluders, so that more shapes should be completed as one compared to the wide condition. (2) Symmetry was manipulated by mirroring one half of the occluded shape (unfamiliar shapes) or picking naturally symmetric animals like butterflies (familiar shapes). Two possible competing hypotheses were formulated for the effect of symmetry: The grouping hypothesis posits that symmetric shapes will be more likely to be completed as one, whilst the axes-reinforcement hypothesis predicted the opposite. (3) Familiarity was manipulated by presenting novel naturalistic-looking stimuli (unfamiliar), or real animal silhouettes (familiar). We hypothesized familiarity to interact with distance, so that more familiar shapes should be completed as two in the wide occluder condition.

### 4.1 Biases in human shape completion

#### Occluder size

The results of the first experiment showed that the distance between the visible parts of a stimulus significantly affected the perception of shapes as one, or two shapes. In the narrow occluder condition, nearly all shapes were completed as one continuous shape, demonstrating the importance of good continuation. In the wide occluder condition, significantly more shapes were completed as two than in the narrow occluder conditions (**Figure 4A**) - suggesting that interpolation becomes more difficult with increasing distance, which is diminishing the role of good continuation. Still, note that even in the wide occluder condition more shapes were completed as one shape than as two shapes, illustrating the fundamental contribution of good continuation.

#### Symmetry

Symmetric stimuli across conditions were shown to be significantly more likely to be completed as two shapes than asymmetric shapes (**Figure 4C**). This is more in line with the axes-reinforcement hypothesis than the grouping hypothesis. The axes-reinforcement hypothesis was based on the finding that the presence of a global axis of symmetry increases the sensitivity to potential local axes of symmetry (Nucci & Wagemans, 2007). To further explore this relationship between local and global symmetry in the current dataset, we asked independent raters to identify local and global axes of symmetry in each stimulus completed as two objects. The results showed that a significant portion of those drawings had both global and local axes of symmetry. Thus, when a global axis of symmetry is present in the original stimulus, observers may become more sensitive to potential local axes of symmetry in their completions. This interpretation is also supported by our finding that asymmetric stimuli were almost never completed with local axes of symmetry, suggesting that without the increased sensitivity induced by global symmetry, observers do not spontaneously impose local symmetry on their completions.

#### Familiarity

Significantly more stimuli were completed as two objects in the familiar condition than in the unfamiliar condition (**Figure 4B**). As hypothesized, this was the case in the wide occluder condition, while no effect was observed in the narrow occluder condition (**Figure 4D**).

#### Summary

Altogether, our results suggest that in a free drawing task with naturalistic stimuli, good continuation is exerting the strongest bias on completions. For the specific cases we studied here this led to significantly more completions of shapes as one rather than two, across all conditions. However, when the distance between visible parts increases and interpolation becomes more difficult, other factors become important. For example, global symmetry seems to increase the likelihood of completing shapes as two, possibly because it increases the visual system’s sensitivity to potential local axes of symmetry (Nucci & Wagemans, 2007). Also, familiarity becomes important, when good continuation violates our previous knowledge about valid completions.

These findings align well with and extend previous research on amodal completion. Our results corroborate the foundational role of good continuation identified by Kanizsa and colleagues (1975, 1979), who demonstrated that this low-level cue can span substantial gaps between visible contours. The persistent tendency to complete shapes as one object even with wide occluders confirms that good continuation operates as a powerful default mechanism in human vision. Similarly, our symmetry findings resonate with Nucci and Wagemans’ (2007) framework: rather than simply grouping symmetric parts together, the visual system appears to use global symmetry as a signal to search for multiple local symmetry axes, thereby increasing the likelihood of two-object completions. Finally, our familiarity effects closely mirror those reported by Yun et al. (2018), who found that object-specific knowledge can override good continuation when it would produce implausible shapes. By demonstrating this effect with complex naturalistic shapes in a free drawing task, we show that the interaction between low-level and higher-level cues generalizes beyond previously used stimuli and tasks.

The systematic pattern of results observed in Experiment 1 demonstrate that human amodal completion involves sophisticated integration of multiple cues operating at different levels of processing. Good continuation serves as the dominant organizing principle, but its influence is modulated by both global symmetry and semantic knowledge. This hierarchical integration, where low-level perceptual features interact with higher-level conceptual knowledge, raises an intriguing question for contemporary AI: do large multimodal models, trained on vast corpora of natural images, exhibit similar completion biases? Experiment 2 examined whether three state-of-the-art LMMs would replicate the human patterns when completing identical stimuli.

### 4.2 Biases in LLM shape completion

Our second experiment aimed to compare human biases in shape completion to those of recent AI models. To this end three publicly available Large Multimodal Models were tasked with completing the same stimuli as our human participants and their results were analyzed equivalent to the human’s drawings. An initial inspection revealed that a large portion of the model outputs failed to meet the task demands, for example, by showing odd additions. To still be able to test shape-completion biases rather than the ability to follow instructions, we only analyzed further the 20 images in each condition that were rated by independent raters as most compliant with the instructions.

The best way to interpret the model output is with respect to their training regimes. In AI, there has long been a distinction between connectionism and symbolism (Xiong et al., 2024): (1) Connectionism uses large neural networks and algorithms to recognize patterns in large datasets (Lecun et al., 2015), which can be seen as analogous to human perception and processing of stimuli (Xiong et al., 2024). (2) Symbolism on the other hand is concerned with rules and symbols, and can be seen as analogous to human reasoning (Newell & Simon, 2007). Notably, the recent leaps in the field driven by the publishing of large public models (e.g. by OpenAI and Google) are based on scaling up connectionist models (Wu et al., 2025). This is also the case for the LMMs we tested in the current study; they were trained on huge datasets, without explicit rules or symbols, to extract feature patterns. Although proponents of connectionism argue reasoning abilities can be achieved by scaling the dataset, and the amount of parameters, the lack of symbolism would suggest that these models operate mainly on a level more analogous to low-level processing than high level reasoning (Sharma, 2024).

#### Occluder Size

Like humans, models completed significantly more shapes as two in the wide occluder condition than in the narrow condition, suggesting that both find it easier to interpolate over small distances than over large distances. This supports the notion that biases towards good continuation might be present in low-level processing.

#### Symmetry

In line with what we know about their training regime, the similarities between humans and models is less evident for higher level properties like symmetry. Although the effect of symmetry trended in the same direction as the human data, it failed to reach significance. More critically, when we examined how models responded to globally symmetric configurations, a key difference emerged: while humans spontaneously produced locally symmetric completions when presented with two drawings that shared a global axis of symmetry, models did not. This differential sensitivity to global symmetry reinforces that humans perceive and represent hierarchical geometric structure beyond low-level local features, whereas models lack this capacity. Rather, they appear to process symmetry (when they do) at a purely local level, failing to extract the hierarchical relationship between global and local symmetry axes that humans readily perceive.

#### Familiarity

With familiar shapes, the models showed significantly less completions as two compared to with unfamiliar shapes in the wide occluder condition - an effect opposite to that observed in humans. This reversal is particularly revealing: humans tended to complete familiar objects as two shapes more often, suggesting they rely on stored global shape priors and semantic knowledge. In contrast, models used two shapes more often for unfamiliar objects. The fact that models show different completion patterns for familiar versus unfamiliar shapes indicates they have learned *some* statistical regularities that distinguish these stimulus types during training. However, the nature of this learning appears fundamentally different from human semantic knowledge. For familiar shapes, models may have encountered similar visual patterns during training that bias them toward single-shape completions, perhaps because these familiar objects frequently appear as unified wholes in the training images. For unfamiliar shapes, lacking such learned patterns, models tend to see more spatially separated contours as spatially distinct entities. Crucially, this pattern is opposite of what semantic, hierarchical knowledge would predict: models are not breaking down familiar multi-part objects into their essential components like humans do, but rather have a tendency to “see” holistic completions. This suggests that whatever regularities models have extracted from their training data, their visual diets likely lead to different representations than the kind of structured, compositional shape representations that allow humans to decompose familiar objects into meaningful parts. This supports the idea that the effect of familiarity, or previous knowledge, relies on a higher level of cognition than the effects of strictly structural features. One possible explanation is that familiarity does not operate as semantic knowledge in the models, because they may not recognize the familiar shapes as semantically meaningful objects. This is in line with prior work showing that convolutional neural networks do not use shape as a primary feature for object classification (Baker et al., 2018). Since shape was the only available feature in our experiment, the models may therefore not have been constrained by semantic knowledge in their completions. However, given the substantial differences in both architecture and task objectives between object classification networks and image-editing generative models, this explanation remains speculative.

Although we did not further analyze the non-compliant outputs, it is worth mentioning that when the models did not meet task demands, they did seem to integrate higher level knowledge in a subset of the outputs (for illustration, see noncompliant outputs in **Figure 6D-I**). This introduces an important point of nuance to these findings. It might be the case that the use of higher level cues is possible for these models, but that integrating them under our task demands, whilst unproblematic for humans, raises an issue for the models. Since our study focussed specifically on the models’ ability to mimic humans performance on a purely shape-based generative shape completion task these results fall outside the current scope, but future studies could investigate how specific task demands, and instruction-following abilities interact with the abilities of multimodal models to integrate lower and higher level cues.

#### Summary

Our findings align with and extend contemporary research on how LMMs process visual information. Previous work has demonstrated systematic divergences between neural networks and human perception, for example, the texture bias in object recognition networks, where models rely disproportionately on local texture features rather than global shape information (Geirhos et al., 2018). Similar findings have emerged showing that neural networks develop local rather than global representations for object recognition (Baker and Elder, 2022). While some recent demonstrations have shown superficial agreement between LMMs and human performance on selected examples, such correspondences do not guarantee that models process visual information in human-like ways. Our systematic manipulation of perceptual cues in amodal shape completion provides critical evidence for where and how model processing diverges from human perception. Our results substantiate the view that current LMMs operate primarily through low-level pattern recognition.

Psychologically, the results of this experiment suggests that a lack of symbolic knowledge is detrimental for amodal shape completion of familiar objects which indeed requires higher order cognition. Given the fact that most natural scenes will involve mainly shapes for which we have a semantic representation (think of a *field*, with *trees*, a *barn* and a *cow* for example) this is a highly relevant aspect of shape completion. The model outputs indicate that purely statistical pattern learning, without access to symbolic or semantic representations, s is not enough to achieve human-level completion performance.

Our results show that large connectionist models are not able to accurately model human performance when it comes to amodal shape completion at this level of abstraction, and under these specific conditions. We speculate that the inclusion of explicit symbolism (e.g., representations of objects as compositions of meaningful parts and relations between them) would model higher level cognition better, and thereby increase performance on this task.

### 4.3 Limitations and future studies

Including the AI models in Experiment 2 introduced several limitations. Because a successful completion inherently requires a good understanding of the task, multiple prompts were tested. Although the final prompts closely matched the instructions given to humans, many model-generated outputs were still non-compliant given the prompt. Attempts at resolving this issue through filtering (based on human raters’ judgements of instruction compliance) resulted in significant data-loss, which decreased the statistical power of Experiment 2. Additionally, even in the remaining datapoints, the model periodically showed completions that would have been impossible for humans to make, such as altering the spatial relationships between visible parts by moving them closer together or further apart. Taken together, this suggests that some of the variability in the model-data reflects the (in)ability of following instructions rather than differences in amodal completion per se. However, given the stringency of the filtering criteria, we do not expect this to be a fundamental challenge to our conclusions.

Interaction with the AI models necessarily relied on prompting, which we aimed to align as closely as possible with the instructions given to human participants while remaining effective for the models. Several considerations regarding prompt use are therefore relevant. First, unlike with human participants, it is not possible to verify whether a model has understood the task beyond inspecting its outputs. Although this could in principle be addressed by using a more conversational, multi-turn interaction, we opted for a single prompt. This decision was motivated both by the desire to ensure consistency across the three AI models tested, and by prior evidence suggesting that language models often perform better with single prompts than with multi-turn conversations (e.g., Laban et al., 2025). Second, the specific wording of the prompt, while closely matching the human instructions, may still introduce subtle biases in model behavior. For instance, instructing the model to “complete the silhouette,” although not explicitly implying a single shape, could nevertheless bias completions toward unified interpretations through the use of the singular form. During prompt development, we tested multiple prompt variants and qualitatively assessed their effects on model outputs. These tests suggested that minor changes in wording or visual properties (e.g., specifying a gray rather than a red occluder, which appeared to improve instruction following) did not meaningfully alter the pattern of results, but primarily affected how reliably the models adhered to the task constraints. For this reason, we adopted a single prompt across all models, not because it was uniquely optimal, but to maintain comparability and control across model conditions.

A final limitation concerns the speed of development in the field of AI. Even though the models used in this study represented the state of the art when data was collected, at the time of writing google has already made advances and published a new version of their image generation LMM (nano banana pro). Preliminary testing suggests that his new version is marginally better at following instructions but still displays many of the same patterns described in this study. Future studies should persist in evaluating the performance of current and future models, and our work can provide a methodological framework to do so.

### 4.4 Conclusion

This study investigated how humans and large multimodal models complete partially occluded shapes as one or two, examining the integration of low-level perceptual cues and higher level semantic knowledge. Our findings reveal fundamental differences in how biological and artificial systems resolve visual ambiguity.

Our results suggest that human amodal completion is guided by a hierarchy of cues. Good continuation serves as the dominant organizing principles under all our experimental conditions. However, when occluders are wide and contour extrapolation becomes more difficult, higher level cues are increasingly important. Global symmetry increases two-shape completions, possibly by heightening sensitivity to potential axes of local symmetry. Familiarity also promotes two-shape completion when single shape completions would yield semantically implausible objects. These findings demonstrate that humans flexibly integrate perceptual principles with semantic knowledge based on task demands.

In contrast, state-of-the-art multimodal AI models showed human-like behaviour only for the low-level cue of occluder size (good continuation). They systematically diverged from human data when higher level reasoning was required, specifically configural global shape features and semantic knowledge. This extends earlier findings for classification models (Baker & Elder, 2022) to generative models for an amodal completion task. The model’s inability to consistently follow instructions further underscores the gap between current AI systems and human cognitive flexibility.

Together, we show that human amodal completion cannot be reduced to a set of fixed perceptual rules operating independently. Rather, the visual system dynamically integrates multiple sources of information in a context-dependent manner. When good continuation provides strong alignment cues, it dominates completion judgments nearly uniformly. However, when contour extrapolation becomes difficult, higher-level knowledge systematically modulates perception. This reveals a visual system that actively uses different perceptual and semantic cues to resolve visual ambiguity.

## Supporting information

Supplementary Matrials

## Acknowledgments

This research was supported by the Deutsche Forschungsgemeinschaft (DFG, German Research Foundation—project number 222641018—SFB/TRR 135 TP C1), the Sectorplan SSH-Breed from the Dutch government, and internal stimulation funds from Erasmus University Rotterdam allocated to the Brain and Cognition team.

## Supplementary material

### S1

#### Screening sheet

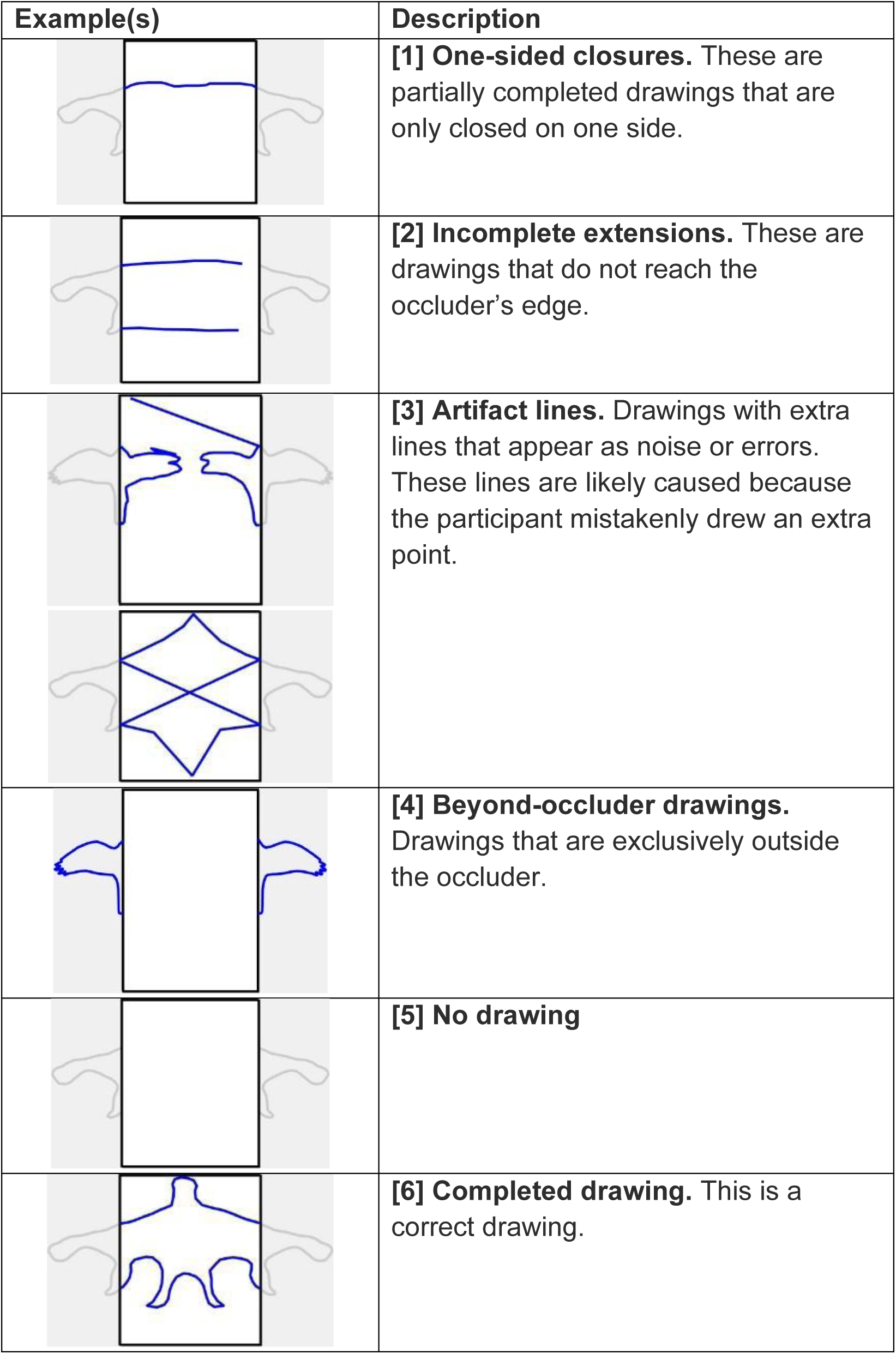

### S2 Global vs Local Symmetry Instructions

In the following task you will be asked to distinguish between global and local axes of symmetry:

**Global Symmetry**

Global symmetry refers to symmetry of an entire array or pattern. A globally symmetric array can be divided by an axis (an imaginary line) where one half is a mirror reflection of the other half. The symmetry encompasses the complete array (left example in the figure below).

For example, a human face has approximate global symmetry along a vertical axis.

**Local Symmetry**

Local symmetry refers to symmetry in just a part of an array or pattern. It creates "pockets" of symmetry within a larger array or pattern: isolated areas that are mirror symmetric (center example in the figure below).

For example, a row of different houses may not be globally symmetric, but each single house might be (locally) symmetric.

Importantly, note that global axes of symmetry and local axes of symmetry can also appear in the same array and are not mutually exclusive (right example in the figure below).

**Examples of different axes of symmetry**

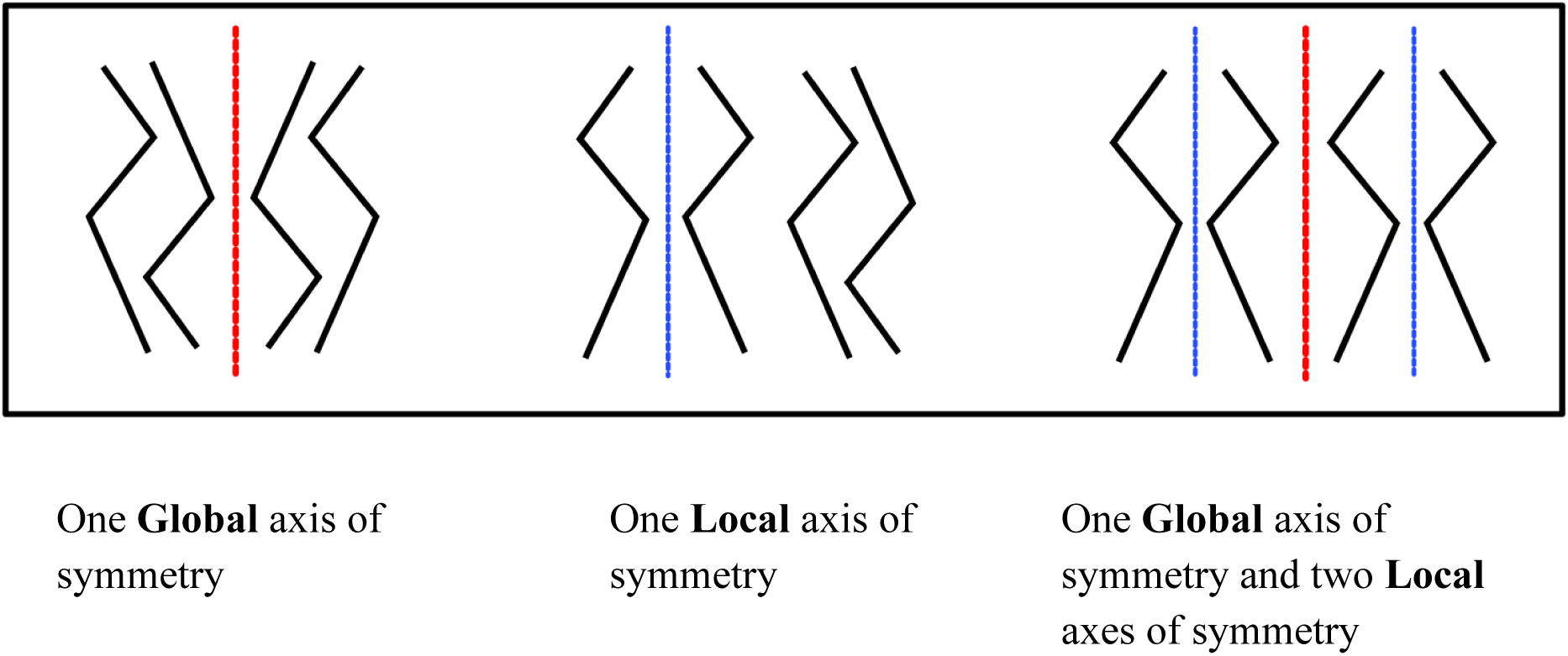

**Your Task:**

In this task, you will be shown arrays containing two hand-drawn shapes. Since these are created by people, focus on identifying where the drawer intended to create symmetry, even if the execution isn’t perfectly symmetrical.

For each array:

1. Examine both drawings carefully
2. And determine:
  - If there is intended global symmetry (affecting the entire array of two drawings)
  - If there are any intended local symmetries (affecting only parts of the array, i.e., single drawings).
3. There can only be a maximum of one global axis of symmetry. However, if you identified axes of local symmetry, indicate for each axis whether it is located to the right or the left of the array’s center

Remember that these are hand-drawn shapes, so look for the intention behind the drawing rather than perfect geometric symmetry.

Please take your time with each image and be specific about which symmetries you observe in each shape.

### S3 - Model Completions as Two - Local and Global symmetry

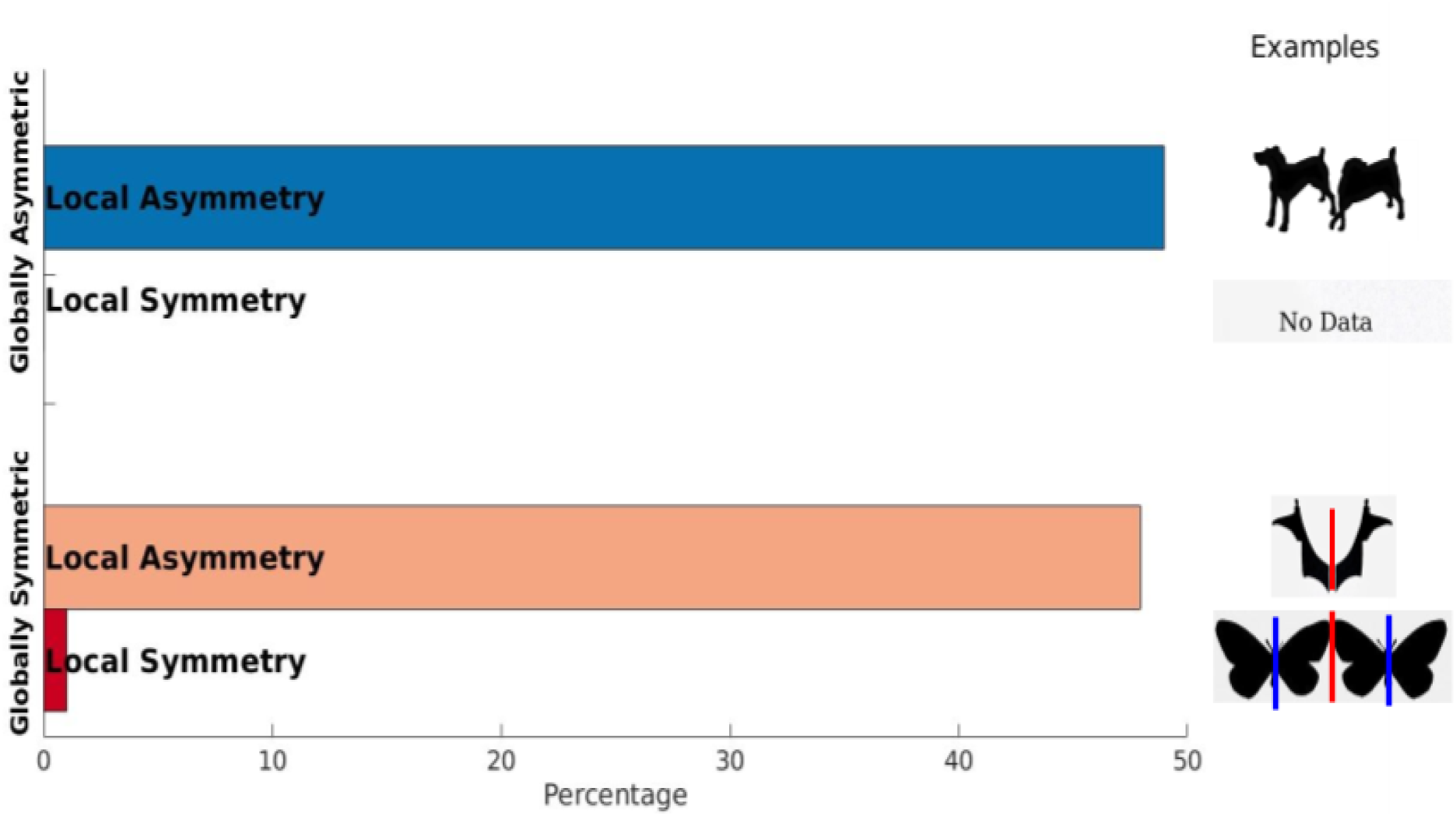

No quality threshold rating was used for this rating task, such that all of the original model outputs categorised as two were analysed in this analysis. 98 out of the total 600 images were completed as two and. Most notably, LMM outputs do lack locally symmetric completions, whereas humans would create local axes of symmetry in almost 50% of their drawings of two shapes. The low occurrences of local symmetry violate the assumptions of a one-sample chi-square test, so a binomial test was performed. The test demonstrated a significant effect in the opposite direction as in human completions, with the models showing a lower probability of using globally symmetric cues to infer locally symmetric shapes (Z = -6.74, p < .001).

