## Supplementary Matrials for "Human and generative AI integrate visual cues differently in a shape completion task"

Supplementary material

**S1**

Screening sheet

| **Example(s)** | **Description** |
|---|---|
| 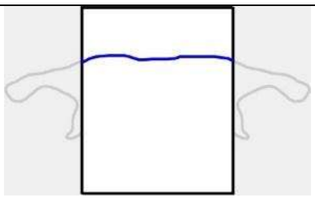 | **[1] One-sided closures.** These are partially completed drawings that are only closed on one side. |
| 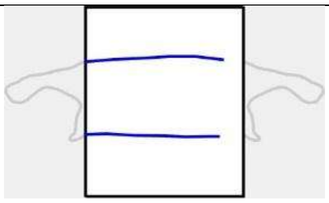 | **[2] Incomplete extensions.** These are drawings that do not reach the occluder’s edge. |
| 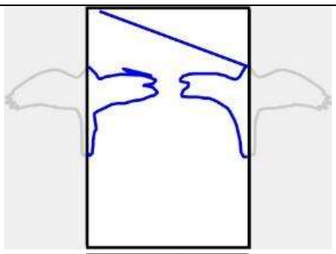 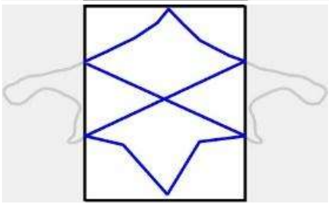 | **[3] Artifact lines.** Drawings with extra lines that appear as noise or errors. These lines are likely caused because the participant mistakenly drew an extra point. |
| 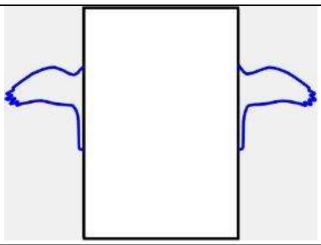 | **[4] Beyond-occluder drawings.** Drawings that are exclusively outside the occluder. |
| 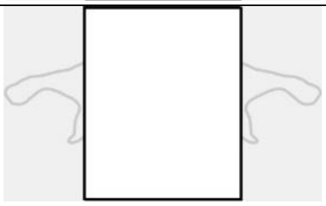 | **[5] No drawing** |
| 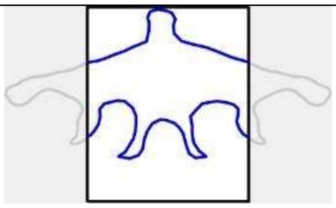 | **[6] Completed drawing.** This is a correct drawing. |

### S2 Global vs Local Symmetry Instructions

In the following task you will be asked to distinguish between global and local axes of symmetry:

#### Global Symmetry

Global symmetry refers to symmetry of an entire array or pattern. A globally symmetric array can be divided by an axis (an imaginary line) where one half is a mirror reflection of the other half. The symmetry encompasses the complete array (left example in the figure below).

For example, a human face has approximate global symmetry along a vertical axis.

#### Local Symmetry

Local symmetry refers to symmetry in just a part of an array or pattern. It creates "pockets" of symmetry within a larger array or pattern: isolated areas that are mirror symmetric (center example in the figure below).

For example, a row of different houses may not be globally symmetric, but each single house might be (locally) symmetric.

Importantly, note that global axes of symmetry and local axes of symmetry can also appear in the same array and are not mutually exclusive (right example in the figure below).

#### Examples of different axes of symmetry

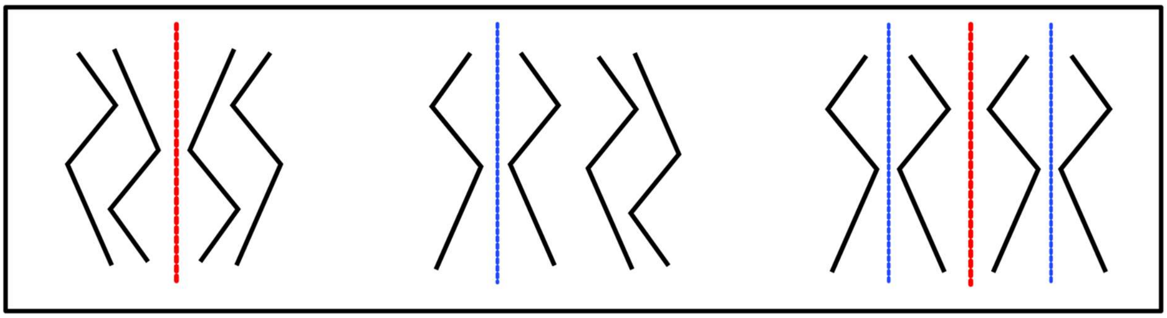

One **Global** axis of symmetry

One **Local** axis of symmetry

One **Global** axis of symmetry and two **Local** axes of symmetry

**Your Task:**

In this task, you will be shown arrays containing two hand-drawn shapes. Since these are created by people, focus on identifying where the drawer intended to create symmetry, even if the execution isn't perfectly symmetrical.

For each array:

1. Examine both drawings carefully
2. And determine:
    - If there is intended global symmetry (affecting the entire array of two drawings)
    - If there are any intended local symmetries (affecting only parts of the array, i.e., single drawings).
3. There can only be a maximum of one global axis of symmetry. However, if you identified axes of local symmetry, indicate for each axis whether it is located to the right or the left of the array's center

Remember that these are hand-drawn shapes, so look for the intention behind the drawing rather than perfect geometric symmetry.

Please take your time with each image and be specific about which symmetries you observe in each shape.

#### S3 - Model Completions as Two - Local and Global symmetry

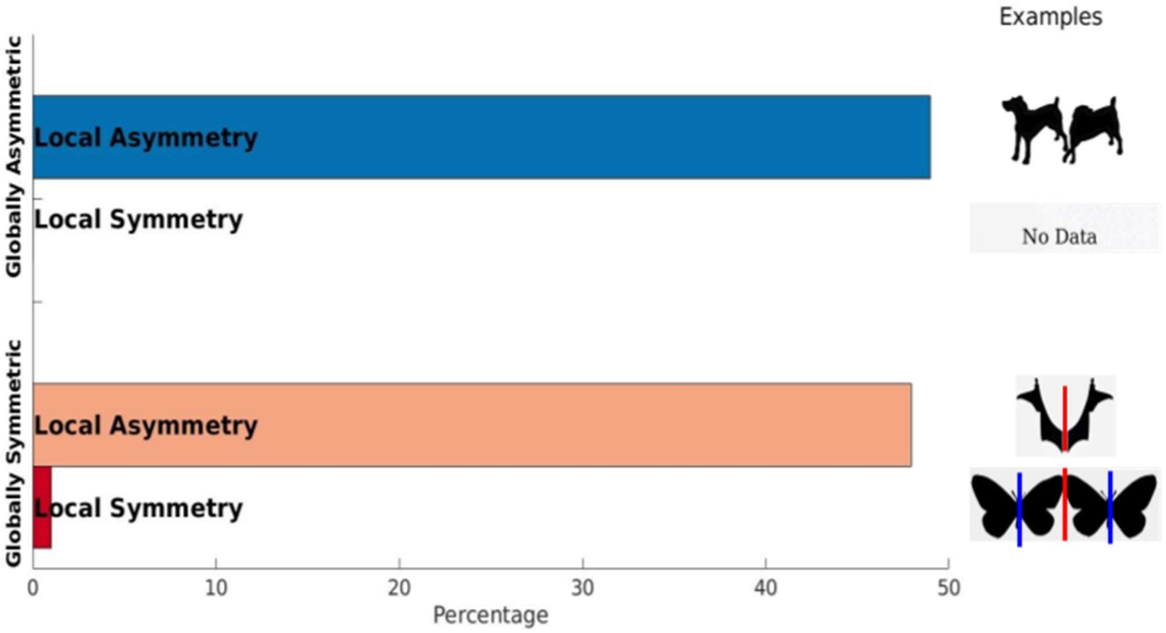


No quality threshold rating was used for this rating task, such that all of the original model outputs categorised as two were analysed in this analysis. 98 out of the total 600 images were completed as two and . Most notably, LMM outputs do lack locally symmetric completions, whereas humans would create local axes of symmetry in almost 50% of their drawings of two shapes. The low occurrences of local symmetry violate the assumptions of a one-sample chi-square test, so a binomial test was performed. The test demonstrated a significant effect in the opposite direction as in human completions, with the models showing a lower probability of using globally symmetric cues to infer locally symmetric shapes ($Z = -6.74$, $p < .001$).
